# Interpretable spatial ecological dynamics reveal ecological retention escape and amplification in pediatric leukemia

**DOI:** 10.64898/2026.07.27.741112

**Authors:** Seung-Hwan Kim

## Abstract

Spatial transcriptomics enables tumor ecosystems to be examined within their native tissue architecture, yet many computational approaches remain primarily descriptive and provide limited insight into how spatial organization may constrain or facilitate disease persistence. Here, we develop a therapy-aware spatial ecological framework that integrates ecological-context discovery, ecological opportunity landscapes, Ornstein–Uhlenbeck (OU)-like retention, Lévy-like escape, and branching-like amplification to characterize pediatric leukemia tissues. Using normal bone marrow reference programs and pediatric leukemia spatial transcriptomic sections, we identify five recurrent candidate ecological contexts with distinct malignant, immune, stromal, vascular, inflammatory, and stress-associated features. These contexts form spatially heterogeneous ecological opportunity landscapes across bone marrow and extramedullary samples. OU-like summaries quantify context-specific attractor position and ecological retention, whereas Lévy-like analyses identify rare cross-context displacements consistent with discontinuous ecological remodeling. Branching-like amplification integrates ecological abundance, latent displacement, opportunity, escape, and escaped-state fraction into an interpretable index of ecological expansion. An independent treatment-associated cohort shows that a primitive-state retention proxy, escape, and amplification generate coherent ecological profiles qualitatively consistent with residual persistence and relapse-associated expansion. Because the available spatial datasets are cross-sectional, these quantities are interpreted as operational summaries of tissue organization rather than direct estimates of temporal evolutionary processes. Together, the framework moves spatial transcriptomic analysis beyond static tissue mapping by providing an interpretable representation of ecological constraint, rare displacement, amplification, and therapy-associated persistence in pediatric leukemia.

## INTRODUCTION

Spatial transcriptomics has transformed tissue biology by enabling gene expression to be interpreted within native spatial architecture (Ståhl et al. 2016; Rao et al. 2021). Unlike bulk sequencing or dissociated single-cell profiling, spatial approaches preserve relationships among malignant, immune, stromal, and vascular compartments within intact microenvironments (Rao et al. 2021; Elhanani et al. 2023). They have consequently advanced the study of cellular heterogeneity, spatial neighborhoods, and tissue organization across developmental, inflammatory, and neoplastic settings (Ji et al. 2020; Lohoff et al. 2022; Vickovic et al. 2022). In cancer, these studies show that progression is shaped not only by malignant-cell-intrinsic programs but also by interactions with immune, stromal, metabolic, and vascular environments (Ji et al. 2020; Elhanani et al. 2023; Hirz et al. 2023). Spatial ecological organization may therefore be an important component of cancer evolution and therapeutic response.

Most computational approaches identify spatial domains, cellular neighborhoods, graph-based dependencies, tissue structures, or cell–cell communication networks (Dries et al. 2021; Hu et al. 2021; Fischer et al. 2023; Cang et al. 2023). Their outputs generally describe locations, co-occurrence patterns, and neighborhood associations of cell states or molecular programs (Rao et al. 2021; Walker et al. 2022; Long et al. 2023), whereas comparatively few methods provide interpretable models of how ecological states constrain, facilitate, or reorganize malignant populations (Walker et al. 2022; Zeng et al. 2022; Elhanani et al. 2023). A related challenge is choosing the biological scale of analysis, because transcript-, cell-, spot-, neighborhood-, and domain-level representations capture different aspects of tissue organization (Atta and Fan 2021; Walker et al. 2022; Fang et al. 2023). Although capture locations provide local measurements, meaningful tissue organization often emerges at an intermediate mesoscale comprising multicellular neighborhoods and tissue compartments (Li and Zhou 2022; Ren et al. 2022; Hu et al. 2024). We previously developed a region-aware bridge-modeling framework that represented spatial transcriptomic architecture at this mesoscale while preserving relationships among gene expression, spatial adjacency, and higher-order tissue structure (Kim 2026b). The present study extends that foundation by asking whether organized tissue states can also be interpreted through an ecological and evolutionary lens.

Tumor evolution can be viewed as an ecological process in which malignant populations occupy, modify, and transition among microenvironmental niches (Merlo et al. 2006; Greaves and Maley 2012; Yang et al. 2014). Under this view, tissue regions are candidate ecological states defined by coordinated malignant, immune, stromal, metabolic, and vascular programs (Ji et al. 2020; Luca et al. 2021; Elhanani et al. 2023). Some states may constrain variation and promote persistence, whereas others may favor expansion, displacement, or treatment resistance (Gatenby and Brown 2020; Vander Velde et al. 2020; Zhang et al. 2024). Therapy can further reshape these landscapes by eliminating sensitive populations, disrupting established interactions, and creating opportunities for residual or resistant states (Gatenby et al. 2009; Obenauf et al. 2015; Rambow et al. 2018). Representations that connect spatial ecological states with constraint, transition, and population-level consequences may therefore provide insight beyond clustering, neighborhood analysis, or tissue segmentation alone (Atta and Fan 2021; Elhanani et al. 2023; Kim 2026b).

Stochastic dynamical models provide quantitative representations of random variation, state transitions, selection, and population expansion during cancer evolution (Beerenwinkel et al. 2015; Altrock et al. 2015; Yin et al. 2019). In our previous iScience study, we developed a hybrid Ornstein–Uhlenbeck (OU)–Branching framework for longitudinal malignant-cell trajectories in pediatric leukemia (Kim 2026a). That framework coupled mean-reverting continuous-state dynamics with stochastic lineage branching and distinguished trajectories dominated by fluctuation, stabilizing constraint, or branching-associated expansion. Together, this longitudinal framework and our region-aware spatial framework provide complementary foundations for the present study (Kim 2026a,b): one models constrained evolution over time, whereas the other represents tissue architecture at an interpretable mesoscale (Kim 2026a; Kim 2026b). Here, we connect these perspectives by developing spatial ecological summaries of retention, opportunity, displacement, and amplification without equating spatial variation with directly observed temporal evolution.

This distinction is necessary because most spatial transcriptomic datasets are cross-sectional snapshots rather than longitudinal observations of the same evolving tissue ecosystem (Atta and Fan 2021; Rao et al. 2021; Moses and Pachter 2022). Nevertheless, cross-sectional organization may contain gradients, transition structures, and state relationships consistent with candidate evolutionary processes when interpreted as spatial summaries rather than temporal trajectories (Pham et al. 2023; Yu et al. 2025). Recurrent contexts may resemble attractor-like states that constrain local variation (Huang et al. 2009; Huang 2013). Rare displacements between compositionally distant contexts may reflect discontinuous ecological remodeling or transitions between state basins (Pisco and Huang 2015; Moris et al. 2016), whereas repeated expansion of favorable states may generate branching-like amplification (Bozic et al. 2010; Greaves and Maley 2012). These concepts motivate OU-like, Lévy-like, and branching-like spatial summaries without implying that formal temporal stochastic processes have been directly observed.

Here, we present a therapy-aware spatial ecological framework that integrates ecological-context discovery, evolutionary opportunity landscapes, OU-like retention, Lévy-like escape, and branching-like amplification. We first identify recurrent pediatric leukemia ecological contexts using interpretable programs derived from normal bone marrow and leukemia-associated features. We then construct context-specific opportunity landscapes and project them onto tissue architecture. OU-like summaries characterize context-specific latent-state position and spatial persistence, whereas Lévy-like analysis identifies rare, large cross-context displacements. Finally, branching-like amplification combines abundance, displacement, opportunity, and escape and is evaluated against independent therapy-associated states.

All quantities are interpreted as operational summaries of spatial ecological organization. OU-like denotes context-specific attraction and persistence, Lévy-like denotes rare and comparatively large cross-context displacement, and branching-like denotes amplification associated with favorable combinations of abundance, opportunity, and escape. By connecting mesoscale spatial architecture with interpretable summaries of ecological constraint, displacement, and amplification, the framework provides a hypothesis-generating bridge among spatial genomics, ecological modeling, and longitudinal evolutionary analysis. Future studies combining spatial, longitudinal, single-cell, clonal, and treatment-response measurements will be needed to determine whether these organizational signatures correspond to directly observed mechanisms of persistence, adaptation, and relapse.

## RESULTS

### A therapy-aware OU–Lévy–Branching framework for spatial leukemia ecology

We first developed a conceptual framework to connect spatial tissue organization with therapy-associated leukemia evolution. In this model, malignant cells occupy spatially structured ecological niches, including hypoxic, stromal-supportive, immune/inflammatory, and vascular regions (Fig. 1A). Local niche structure is represented as an ecological attractor landscape, where OU-like retention summarizes stabilizing forces that constrain malignant or microenvironmental states within spatial basins (Fig. 1B). Therapy is represented conceptually as a perturbation that weakens pre-existing attractors and reshapes the ecological landscape, allowing resistant niches to expand after treatment (Fig. 1C). Within this perturbed landscape, rare discontinuous ecological transitions are represented as Lévy-like escape events (Fig. 1D), followed by branch-mediated amplification of resistant or relapse-associated states (Fig. 1E). Thus, the overall framework links local spatial constraint, rare ecological escape, branching-like expansion, and therapy-associated persistence.

**Figure 1.**
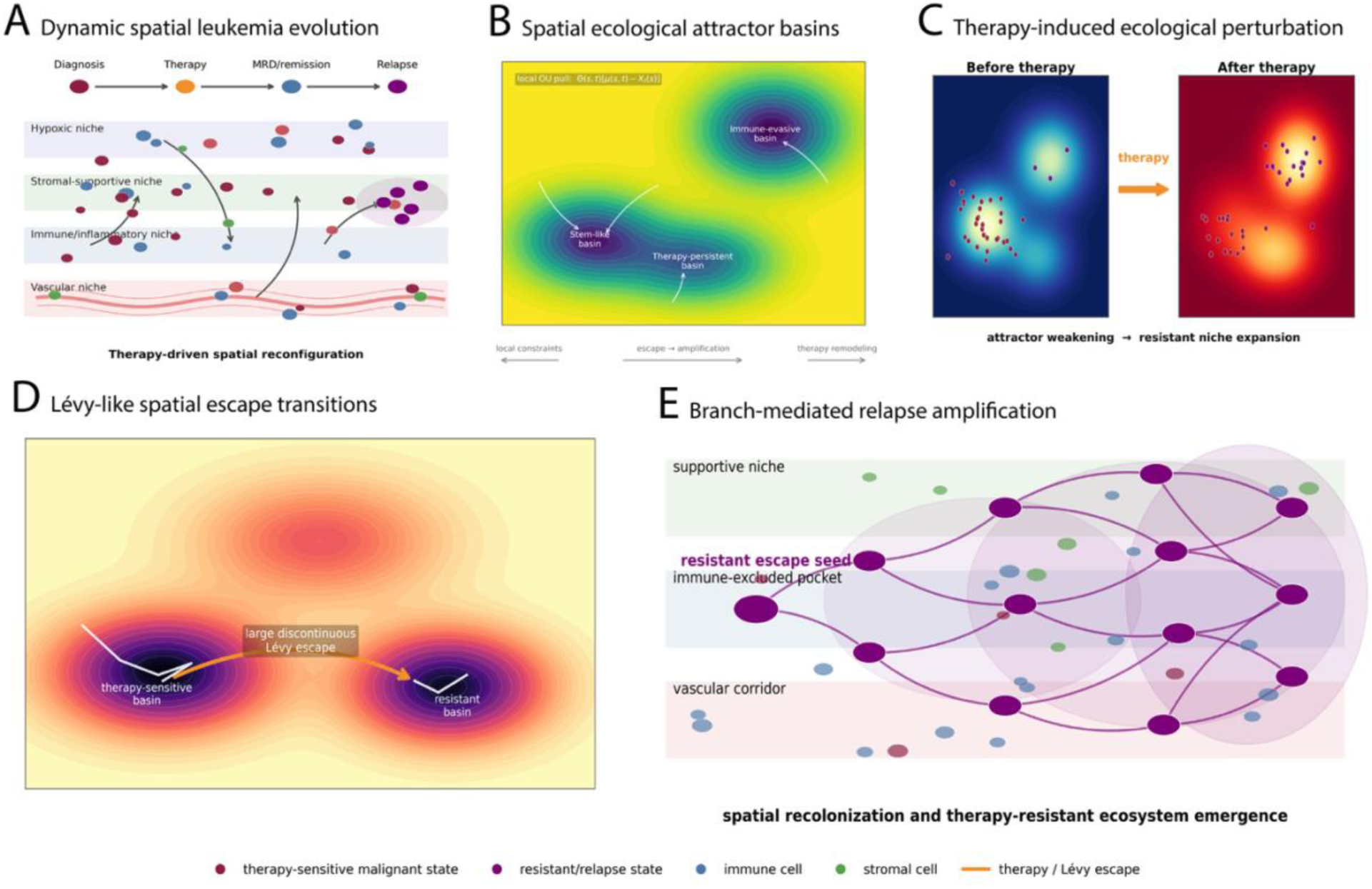
Therapy-aware spatial ecological framework integrating OU-like retention, Lévy-like escape, and branching amplification. Conceptual overview of the proposed therapy-aware spatial ecological framework. **(A)** Spatial ecological organization of pediatric leukemia tissues showing representative ecological niches, including hypoxic, stromal-supportive, immune/inflammatory, and vascular regions. **(B)** Ecological opportunity landscape illustrating attractor-like ecological basins in which local spatial organization is summarized using OU-like retention. **(C)** Therapy perturbs the ecological landscape, weakening existing attractors and creating conditions that permit ecological remodeling and persistence. **(D)** Rare discontinuous ecological transitions are represented as Lévy-like escape events connecting distinct ecological basins. **(E)** Escaped ecological states undergo branching-like amplification, leading to expansion of therapy-persistent or relapse-associated ecological populations. The framework integrates spatial ecological context discovery, opportunity landscapes, OU-like attractor summaries, Lévy-like ecological escape, branching-like amplification, and therapy-associated persistence into a unified ecological representation of pediatric leukemia.

### Spatial transcriptomics identifies reproducible ecological contexts in pediatric leukemia tissues

To define spatial ecological states, we first constructed reference ecological signatures from normal bone marrow samples in GSE253355 (Fig. 2A). These signatures captured hematopoietic, stromal, vascular, immune, inflammatory, and stress-associated programs. We then applied unsupervised ecological context discovery to Visium spatial transcriptomic profiles, identifying five spatial ecological contexts, S1–S5, from the ecological feature matrix (Fig. 2B and C). These contexts showed distinct feature profiles and site biases. S1 was enriched for blast-committed myeloid and hypoxia/stress features; S2 for primitive AML/HSPC-like and ECM-associated features; S3 for monocyte–macrophage, B-cell-associated, and candidate immune-suppressive features; S4 for inflammatory stromal–vascular/ECM features; and S5 for T/NK immune-active features.

**Figure 2.**
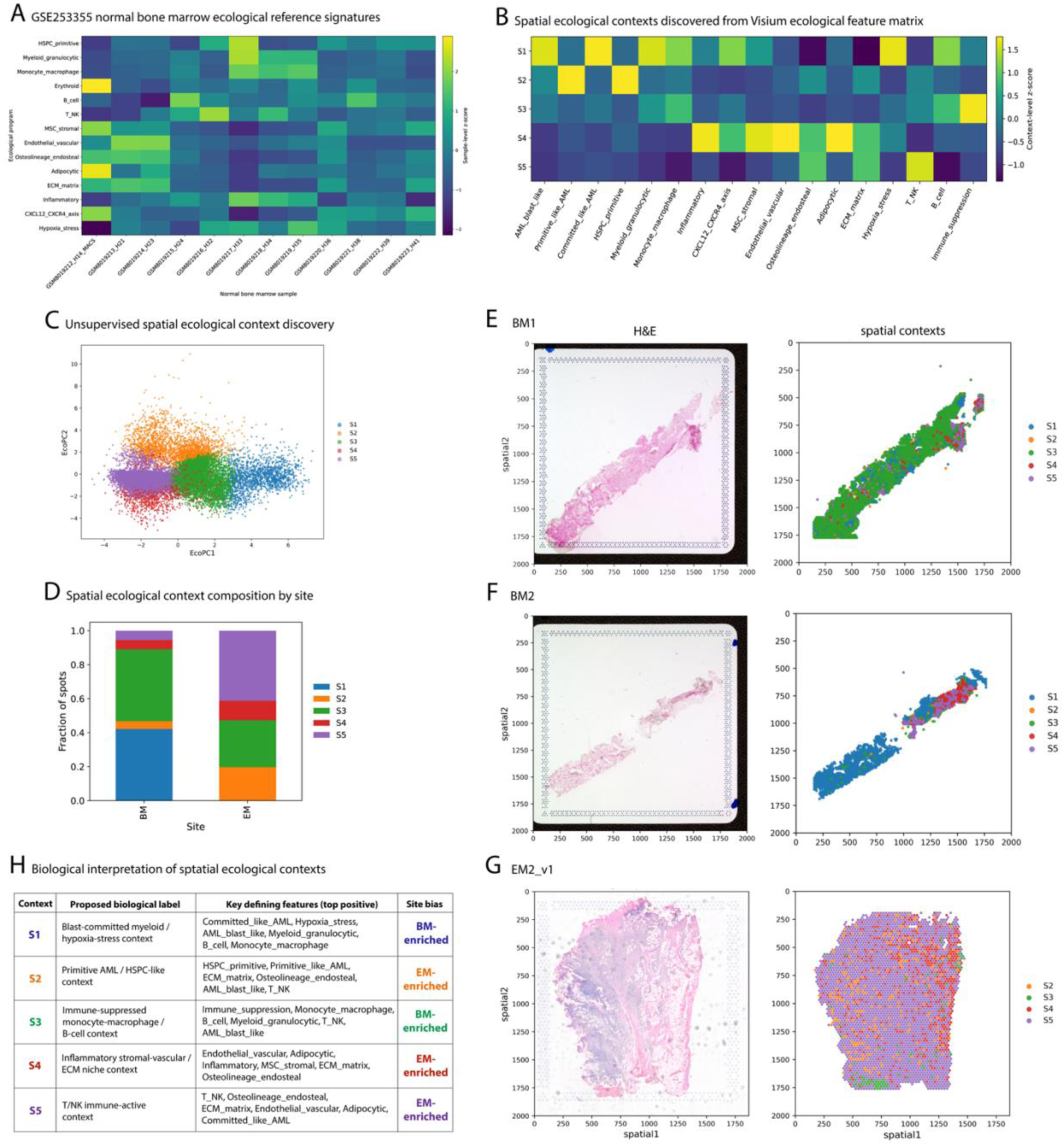
Spatial ecological context discovery identifies reproducible ecological niches in pediatric leukemia. Spatial ecological context discovery from normal bone marrow and pediatric leukemia spatial transcriptomic datasets. **(A)** Reference ecological signatures derived from normal bone marrow samples. **(B)** Context-level standardized ecological feature profiles for the five spatial ecological contexts identified by unsupervised clustering. **(C)** Two-dimensional projection of ecological contexts in latent ecological space. **(D)** Relative abundance of ecological contexts across bone marrow and extramedullary leukemia samples. **(E–G)** Projection of ecological context assignments onto representative Visium tissue sections, demonstrating organized spatial localization of S1–S5 within leukemia tissues. **(H)** Biological interpretation of the five ecological contexts based on their dominant ecological programs. These analyses establish reproducible spatial ecological contexts that serve as the foundation for downstream ecological modeling.

Context composition differed between bone marrow and extramedullary samples (Fig. 2D). Spatial projection onto representative tissue sections showed that these ecological contexts were not randomly distributed but formed organized tissue patterns across BM1, BM2, and EM2_v1 sections (Fig. 2E–G). Together, these results indicate that pediatric leukemia tissues contain reproducible spatial ecological contexts with interpretable biological features and site-specific organization.

### Spatial ecological contexts define an evolutionary opportunity landscape

We next asked whether the discovered ecological contexts could be organized into an evolutionary opportunity landscape. Context-level relationships revealed a structured spatial niche graph in which S1, S2, S3, S4, and S5 occupied distinct positions in ecological space (Fig. 3A). Opportunity scores varied strongly across contexts, with S1 and S4 showing the highest positive opportunity values, whereas S5 showed the lowest opportunity score (Fig. 3B). Within the predefined opportunity-score formulation, the contexts differed in their relative candidate ecological potential for malignant persistence, adaptation, or transition.

**Figure 3.**
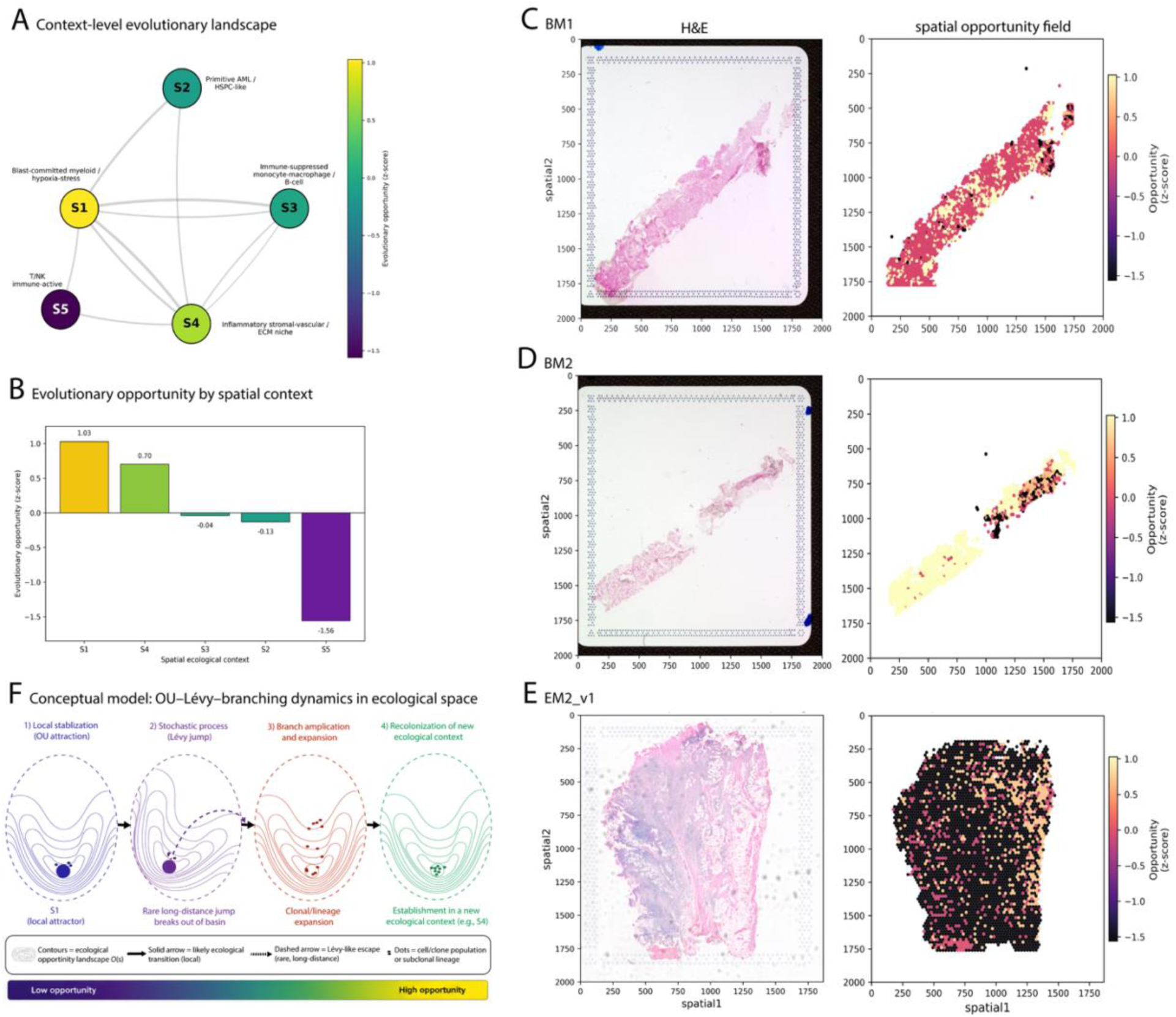
Spatial ecological opportunity landscapes summarize context-specific evolutionary potential. Construction of spatial ecological opportunity landscapes from ecological context organization. **(A)** Graph representation of relationships among the five ecological contexts. **(B)** Context-level opportunity scores summarizing relative ecological potential across S1–S5. **(C–E)** Projection of opportunity scores onto representative bone marrow and extramedullary tissue sections, illustrating heterogeneous spatial opportunity landscapes. **(F)** Conceptual ecological model showing the proposed relationship between local ecological retention, rare ecological escape, branching-like amplification, and recolonization of alternative ecological niches. Opportunity scores provide interpretable summaries of candidate ecological potential rather than direct measurements of evolutionary fitness.

Projection of opportunity scores back onto tissue coordinates showed spatially heterogeneous opportunity fields across representative samples (Fig. 3C–E). BM1 and BM2 contained focal regions of higher opportunity embedded within broader tissue architecture, whereas EM2_v1 showed a more globally altered opportunity pattern. We summarized these relationships in a conceptual OU–Lévy–Branching model in ecological space, in which local stabilization within an attractor can be followed by rare long-distance ecological escape, branch expansion, and recolonization of a new ecological context (Fig. 3F).

### OU-like spatial summaries reveal context-specific ecological attractor structure

We next quantified OU-like spatial summaries across the five ecological contexts using a one-dimensional latent ecological-state axis and an AR(1)-like spatial autocorrelation approximation. Context-specific attractor positions, *μ*, projected onto tissue coordinates revealed marked spatial heterogeneity in the latent ecological state across the representative bone marrow section (Fig. 4A). By contrast, the derived OU-like rate index, *θ* = −log(*ρ*), summarized the rate at which latent-state similarity decayed along the spatial pseudo-order (Fig. 4B). Accordingly, lower *θ* values corresponded to higher positive lag-1 spatial autocorrelation and more persistent local organization, whereas larger *θ* values indicated weaker spatial persistence and more rapid decorrelation.

**Figure 4.**
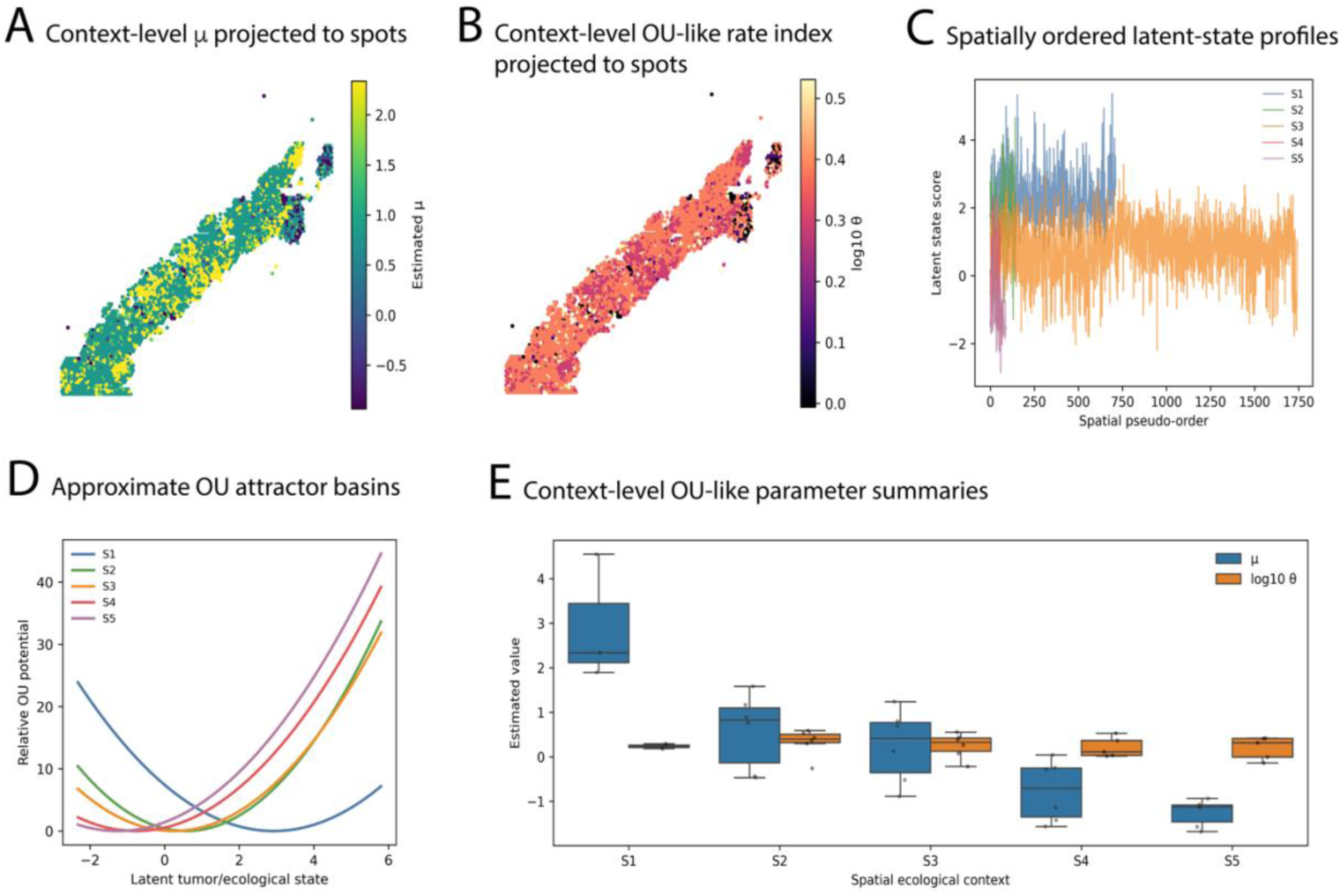
OU-like spatial ecological summaries reveal context-specific attractor organization. Context-level OU-like ecological summaries across spatial ecological contexts. **(A)** Spatial projection of estimated attractor positions (*μ*) onto representative tissue sections. **(B)** Spatial projection of estimated OU-like spatial rate indices (*θ*). **(C)** Spatially ordered latent ecological-state profiles across S1–S5. **(D)** Approximate OU-like ecological attractor basins reflect context-specific rate indices. **(E)** Summary of estimated *μ* and log_10_*θ* values across ecological contexts. Because the analyzed spatial transcriptomic datasets are cross-sectional, *μ* and *θ* are interpreted as statistical summaries of spatial ecological organization rather than direct temporal Ornstein–Uhlenbeck process parameters.

Spatially ordered latent-state profiles further demonstrated that the ecological contexts occupied distinct regions of the latent-state axis and followed different spatial patterns (Fig. 4C). S1 was concentrated at higher latent-state values, consistent with stronger AML- and primitive-like ecological features, whereas S4 and S5 were shifted toward lower latent-state regions. S2 and S3 occupied intermediate positions and showed broader overlap across the spatial ordering. These patterns indicate that the five contexts differ not only in their ecological composition but also in their characteristic latent-state position and degree of spatial continuity.

Approximate OU-like potential curves, constructed from the context-level means of *μ* and the available *θ* estimates, showed distinct attractor-like basin structures across S1–S5 (Fig. 4D). S1 exhibited the most strongly shifted high-*μ* basin, whereas S4 and S5 occupied lower latent-state positions that were clearly separated from the S1-like state. Differences in basin curvature reflected variation in the estimated spatial rate index rather than directly fitted temporal restoring forces. Context-level summaries across samples confirmed substantial differences in both *μ* and log_10_*θ* among the five ecological contexts (Fig. 4E).

Together, these analyses show that recurrent leukemia ecological contexts possess distinguishable latent-state centers and spatial autocorrelation structures. The OU-like summaries therefore provide interpretable descriptors of context-specific attractor position, spatial persistence, and decorrelation. Because these quantities were derived from cross-sectional tissue organization rather than repeated temporal measurements, they should be interpreted as operational summaries of spatial ecological structure rather than direct estimates of temporal Ornstein–Uhlenbeck dynamics.

### Lévy-like ecological escape captures rare cross-context transitions

We next evaluated whether spatial ecological organization contains rare cross-context transitions consistent with Lévy-like escape. The spatial niche landscape showed context relationships in ecological space, with opportunity values mapped onto context nodes and transition edges (Fig. 5A). Cross-context neighbor analysis revealed that several context pairs, particularly S2–S3, S2–S5, and S4–S5, contributed disproportionately to cross-context spatial transitions (Fig. 5B). The ecological jump-distance distribution was right-skewed, and a rare-jump cutoff identified unusually large cross-context transitions (Fig. 5C).

**Figure 5.**
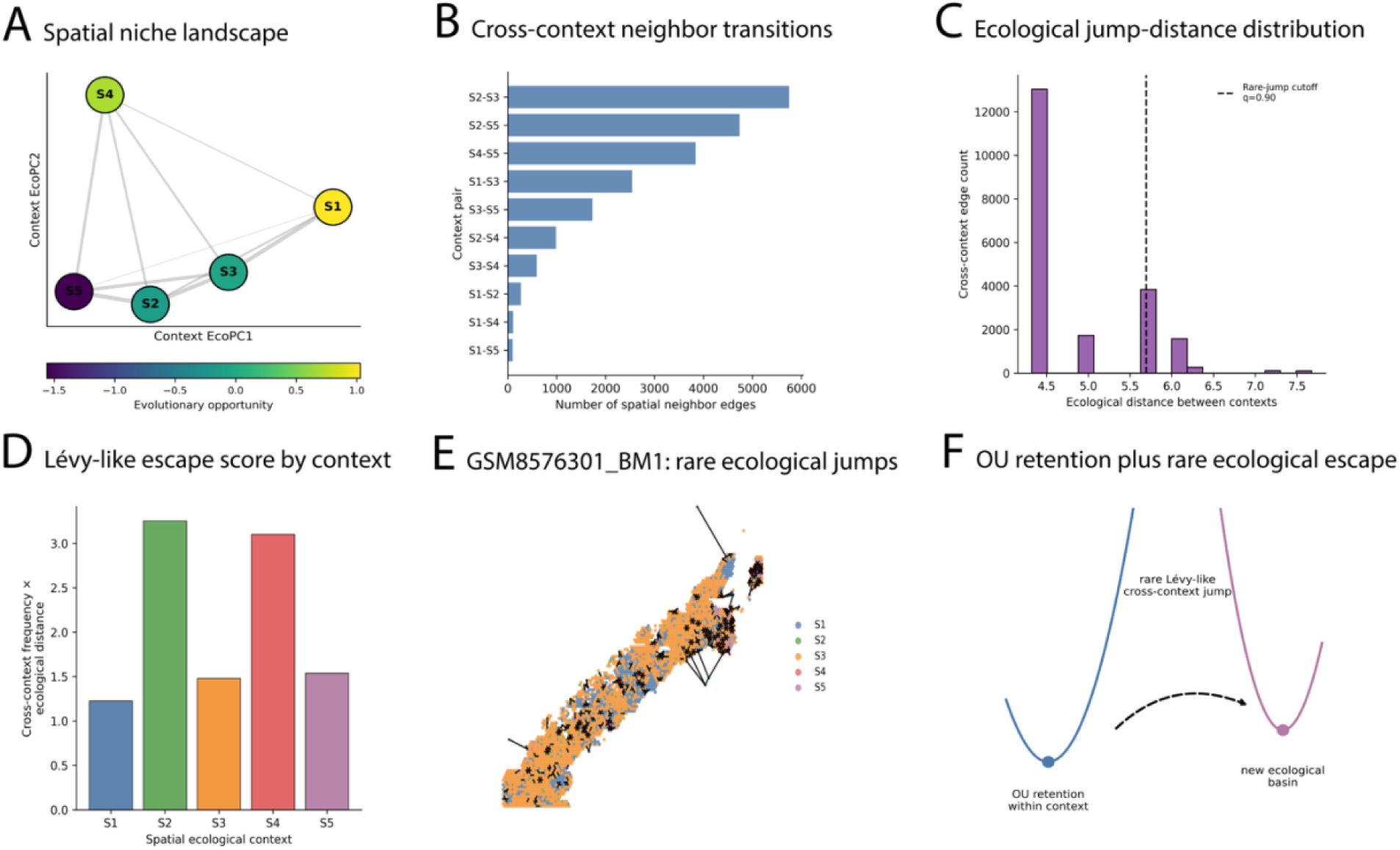
Lévy-like ecological escape identifies rare cross-context spatial transitions. Analysis of rare ecological transitions between spatial ecological contexts. **(A)** Spatial ecological niche network showing context-level relationships and opportunity scores. **(B)** Cross-context neighbor transition matrix summarizing ecological transitions between S1–S5. **(C)** Empirical distribution of ecological jump distances among cross-context spatial-neighbor edges. The dashed line indicates the 90th percentile of the pooled cross-context ecological-distance distribution, which was used as the rare-jump threshold. **(D)** Context-specific Lévy-like escape scores, calculated as the product of the outgoing cross-context edge frequency and the mean ecological distance of those edges. **(E)** Projection of rare cross-context edges, defined as cross-context neighbor relationships with ecological distances at or above the 90th-percentile threshold, onto the representative GSM8576301_BM1 section. **(F)** Conceptual illustration of OU-like ecological retention, Lévy-like rare ecological escape, and subsequent transition into alternative ecological attractor basins. Lévy-like escape is interpreted as a statistical summary of rare discontinuous ecological transitions rather than direct observation of a mathematical Lévy process.

Context-level Lévy-like escape scores differed across ecological contexts, with S2 and S4 showing the highest escape scores (Fig. 5D). Spatial projection of rare ecological jumps in BM1 identified localized regions where rare long-distance transitions occurred within the tissue architecture (Fig. 5E).

Conceptually, these findings support a model in which OU-like retention maintains local ecological structure, but rare Lévy-like cross-context jumps allow cells or ecological states to access distant basins that may support resistant behavior (Fig. 5F).

### Branching amplification links ecological escape to therapy-associated persistence

Finally, we integrated branching-like amplification with therapy-response validation. We modeled the final stage of the framework as a transition from resident OU-retained states to escaped states and then to branching-like amplification (Fig. 6A). Embedding of resident, escaped, and amplified states showed that amplified states formed localized regions within the cell-state landscape rather than being uniformly distributed (Fig. 6B). A state amplification network further identified MonoDC-like states as a dominant amplified node connected to Prog-like, HSC-like, not-malignant, and other-state compartments (Fig. 6C).

**Figure 6.**
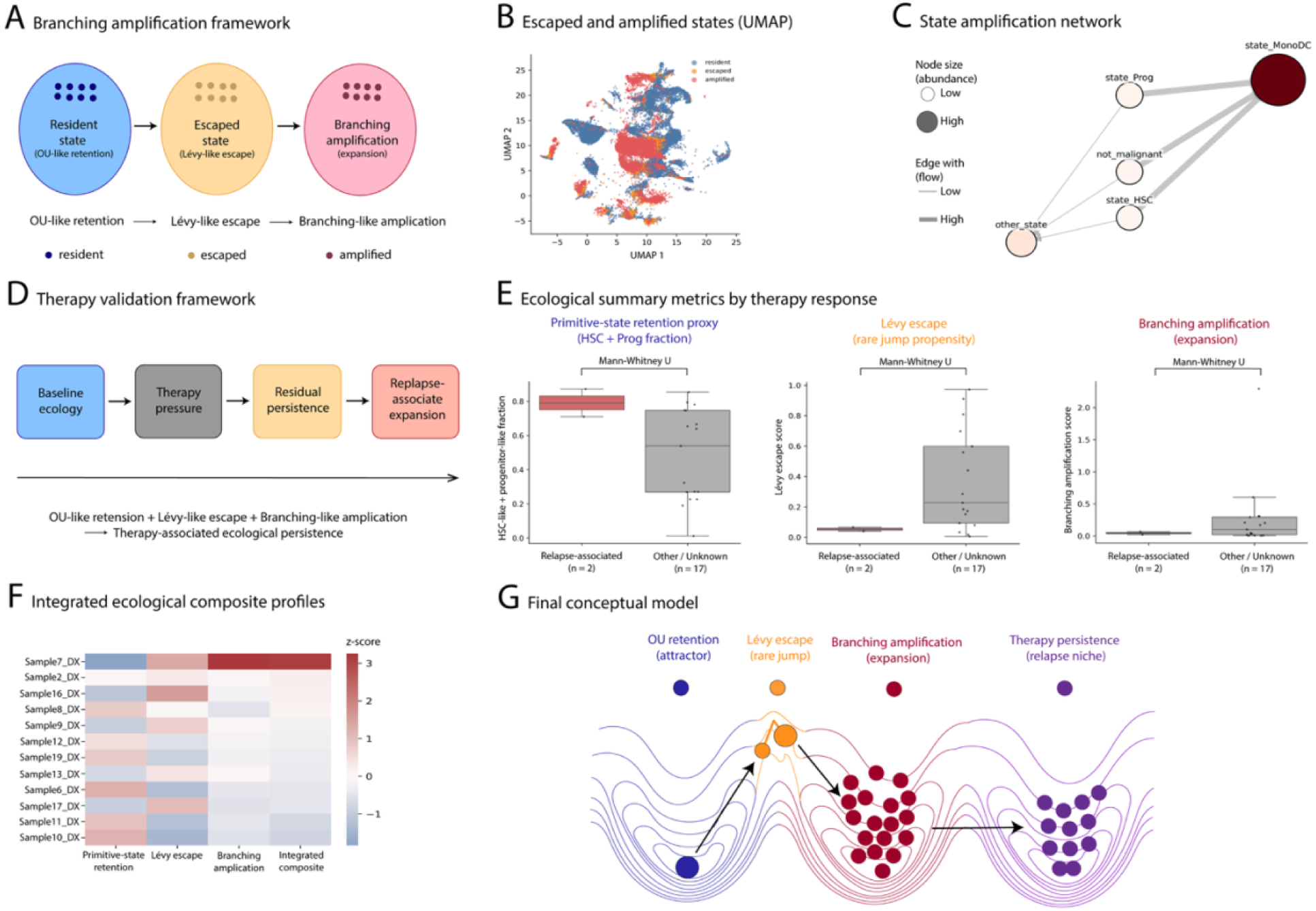
Integrated branching-like amplification and therapy-associated ecological persistence. **(A)** Conceptual progression from an OU-like retained resident state to Lévy-like escape and branching-like amplification. **(B)** Low-dimensional embedding of resident, escaped, and amplified states. **(C)** State-amplification network, with node size proportional to state abundance and edge width proportional to amplification strength. **(D)** Therapy-validation framework linking baseline ecology, therapy pressure, residual persistence, and relapse-associated expansion. **(E)** Primitive-state retention, Lévy-like escape, and branching-like amplification in relapse-associated versus other/unknown samples. Primitive-state retention was defined as the summed HSC-like and progenitor-like fractions and is distinct from the spatial OU-like quantities in Figure 4. Boxes show the interquartile range, center lines the median, whiskers 1.5 times the interquartile range, and points individual samples. Two-sided Mann–Whitney U tests were used; *P* values were unadjusted. Groups contained *n* = 2 relapse-associated and *n* = 17 other/unknown samples and were interpreted exploratorily. **(F)** Integrated heatmap of the three ecological metrics and their summed standardized composite score across diagnosis samples. **(G)** Conceptual synthesis of ecological retention, escape, amplification, and therapy-associated persistence. These analyses represent exploratory biological consistency rather than clinical prediction.

To connect these ecological summaries with therapy response, we organized the validation framework as baseline ecology, therapy pressure, residual persistence, and relapse-associated expansion (Fig. 6D).

Relapse-associated samples showed a higher primitive-state retention proxy, defined as the combined HSC-like and progenitor-like cell fraction, than other or unknown samples, whereas Lévy-like escape and branching-like amplification were more variable in the other or unknown group (Fig. 6E). This compositional retention proxy was used only for treatment-associated validation and was distinct from the spatial *ρ*- and *θ*-based OU-like summaries estimated in Figure 4. Although this validation cohort is limited, the direction of the integrated summaries is consistent with the model that therapy-associated persistence reflects a combination of primitive-state retention, escape potential, and expansion.

An integrated ecological composite heatmap combined the primitive-state retention proxy, Lévy-like escape, branching-like amplification, and the summed standardized composite score across diagnosis samples (Fig. 6F). The samples with the highest composite scores showed coordinated elevation across multiple ecological summary measures, including the sample with the strongest branching-like amplification. We summarized the full model as a sequence of OU-like retention, Lévy-like rare jump, branching amplification, and therapy-persistent relapse niche formation (Fig. 6G). Together, these results support a layered spatial ecological model in which attractor-like retention, rare ecological escape, and branching-like expansion jointly provide interpretable summaries of therapy-associated leukemia persistence.

## DISCUSSION

Spatial transcriptomics enables molecular profiles to be interpreted within native tissue architecture, but many computational approaches remain primarily descriptive, emphasizing clustering, neighborhoods, communication, or graph structure. Here, we present a therapy-aware spatial ecological framework that integrates ecological-context discovery, evolutionary opportunity, OU-like retention, Lévy-like escape, and branching-like amplification in pediatric leukemia. The framework is intended to complement established spatial methods by providing interpretable summaries of ecological persistence, displacement, amplification, and therapy-associated remodeling.

This study builds on two complementary methodological foundations. Our Bioinformatics Advances study established a region-aware mesoscale representation of spatial transcriptomic tissue architecture (Kim 2026b), whereas our iScience study modeled constrained leukemia evolution using longitudinal OU and branching dynamics (Kim 2026a). The present work connects these perspectives by treating recurrent spatial contexts as candidate ecological states and summarizing their retention, opportunity, escape, and amplification. Spatial representation, longitudinal evolutionary inference, and ecological integration therefore form related but distinct components of a broader methodological framework.

Using ecological gene programs informed by normal bone marrow and leukemia biology, we identified five candidate spatial ecological contexts with distinct biological features and tissue distributions. These contexts provided an interpretable intermediate representation between spot-level measurements and higher-level ecological modeling. Rather than defining definitive cell identities, they summarized composite tissue environments, including primitive leukemia, inflammatory stromal–vascular, immune-suppressed, and immune-active states.

The resulting evolutionary opportunity landscape showed that ecological potential was spatially heterogeneous across bone marrow and extramedullary tissues. Contexts enriched for primitive leukemia, stromal support, vascular remodeling, or inflammatory programs generally showed higher opportunity scores, whereas immune-active contexts showed lower values. These scores are not direct measurements of evolutionary fitness; they are composite summaries of ecological programs hypothesized to promote or restrict malignant persistence. Their value lies in representing spatially uneven tissue environments in an interpretable form.

OU-like summaries further distinguished contexts by latent-state position and spatial persistence. This application differs fundamentally from our earlier longitudinal OU analysis (Kim 2026a). Because the present spatial datasets are cross-sectional, the estimated quantities cannot be interpreted as temporal equilibrium positions or mean-reversion rates. Instead, they summarize how strongly spatial ecological states are organized around characteristic context-specific positions. The shared connection between the two studies is conceptual rather than inferential: both use stochastic-dynamical ideas to distinguish constraint, fluctuation, and expansion, but only the longitudinal study directly models temporal trajectories.

Lévy-like escape analysis identified uncommon but comparatively large cross-context displacements. These events may reflect transitions across tissue boundaries, movement between distinct ecological regimes, or spatial remodeling into niches with different opportunity profiles. We do not claim that the data establish a formal Lévy process. Demonstrating such a process would require explicit generative- model comparison, denser sampling, and preferably longitudinal spatial observations. Here, Lévy-like terminology distinguishes frequent local transitions from rare, larger ecological displacements.

Branching-like amplification integrated abundance, displacement, opportunity, escape, and escaped-state fraction into an operational measure of ecological expansion. The score does not represent directly observed cell division, lineage branching, or clonal fitness. Instead, it summarizes candidate states that are both prominent and ecologically positioned for expansion. This construction is conceptually related to the branching component of our longitudinal framework, in which successful malignant lineages expand after escaping stabilizing constraints (Kim 2026a), while remaining distinct from formal branching-process inference.

Therapy-associated validation showed that primitive-state retention, Lévy-like escape, and branching-like amplification produced coherent ecological profiles in an independent cohort. The primitive-state retention proxy represented the combined HSC-like and progenitor-like fraction and was distinct from the spatial ρ- and θ-based OU-like summaries estimated from GSE279576. Samples with stronger combinations of primitive-state retention, escape, and amplification tended to show more relapse-associated profiles. These findings support the biological plausibility of the framework, but the small relapse-associated group precludes clinical interpretation. The analysis should therefore be viewed as an exploratory consistency assessment rather than a validated prediction model.

Several limitations remain. First, all spatial analyses are cross-sectional, so OU-like, Lévy-like, and branching-like quantities are operational summaries rather than direct estimates of temporal evolutionary processes. Second, paired spatial and longitudinal pediatric leukemia datasets are unavailable, preventing direct assessment of whether diagnosis-associated contexts persist or expand during therapy and relapse. Third, ecological-context annotations are based on composite gene programs and should be considered candidate biological interpretations. Fourth, branching-like amplification is not supported by lineage tracing or phylogenetic reconstruction. Fifth, rare-transition estimates depend on spatial resolution, neighborhood definition, and threshold choice. Finally, the therapy-response analysis includes few relapse-associated samples and requires confirmation in larger cohorts.

Future studies combining spatial transcriptomics with longitudinal sampling, clonal barcoding, lineage tracing, single-cell sequencing, and treatment-response measurements could directly test whether attractor-like states predict persistence, rare ecological displacements precede treatment escape, and branching-like amplification corresponds to clonal expansion. Such data would support joint models of spatial organization and temporal malignant-cell evolution within a single probabilistic framework.

The framework may also be applicable to other hematologic malignancies and solid tumors, where ecological heterogeneity and treatment-induced remodeling influence progression. Opportunity landscapes could support comparisons among primary and metastatic sites, treatment-naive and resistant tumors, or immune-active and immune-excluded regions. These ecological summaries could also serve as interpretable inputs to graph neural networks, generative models, or spatial foundation models, preserving biologically meaningful intermediate quantities rather than relying exclusively on opaque latent representations.

In summary, this study presents an interpretable framework that connects spatial ecological contexts with opportunity, OU-like retention, Lévy-like escape, branching-like amplification, and therapy-associated persistence. By integrating mesoscale spatial representation with stochastic-dynamical concepts, the framework moves beyond static tissue mapping and provides a hypothesis-generating basis for studying ecological constraint, displacement, amplification, and adaptation in pediatric leukemia.

## METHODS

### Study design and overview

We developed a therapy-aware spatial ecological modeling framework to summarize leukemia tissue organization through five connected layers: spatial ecological context discovery, evolutionary opportunity scoring, OU-like attractor summaries, Lévy-like ecological escape, and branching-like amplification with therapy-response validation. The analysis used publicly available spatial transcriptomic and single-cell leukemia datasets. The spatial analyses were interpreted as cross-sectional ecological summaries rather than direct longitudinal spatial dynamics. Treatment-associated analyses were used as an independent exploratory biological consistency assessment of the ecological framework.

### Datasets

Normal bone marrow reference signatures were constructed from GSE253355. Spatial leukemia ecological contexts were inferred from Visium spatial transcriptomic sections from GSE279576, including bone marrow and extramedullary leukemia samples. Branching-like amplification and therapy-associated analyses used previously generated GSE235923-derived data, including the labeled secondary-cohort AnnData object, the relapse manifest gse235923_manifest_rel.csv, and the diagnosis-cohort outcome table gse235923_dx_secondary_outcomes.csv. These derivative files, originally generated through reference mapping and label transfer from the labeled GSE235063 primary cohort followed by projection onto the primary ecological backbone, were used as analysis inputs; the upstream annotation procedure was not repeated in the present study. The upstream annotation procedure was not repeated as part of the current workflow. The labeled GSE235923 object and derived outcome table were generated through reference mapping and label transfer from the labeled GSE235063 primary cohort, followed by projection onto the primary ecological backbone. Processed AnnData objects, ecological score matrices, spatial coordinates, inferred context labels, sample manifests, and summary tables were used as inputs for figure generation and downstream modeling.

### Ecological program scoring

For each spatial spot (Visium capture location) or single cell, we computed ecological program scores representing hematopoietic, malignant, immune, stromal, vascular, inflammatory, extracellular matrix (ECM), hypoxia, and stress-associated biological states. Gene-program scores were calculated from curated marker-gene sets using normalized expression values and were standardized within each dataset to facilitate comparison across samples. Standardization was performed using feature-wise z-score normalization,

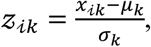

where *x*_i*k*_ is the score of ecological feature *k* for spatial location or cell i, and *μ_k_* and *σ_k_* denote the corresponding dataset-specific mean and standard deviation. The standardized values formed an ecological feature matrix,

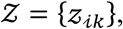

in which rows correspond to spatial locations (or cells) and columns correspond to ecological programs. This standardized ecological feature matrix served as the input for spatial ecological context discovery and all downstream analyses.

To summarize the relative ecological potential of each spatial location, we defined an evolutionary opportunity score as a weighted linear combination of the standardized ecological features,

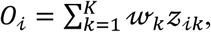

where *O*_i_ is the opportunity score for spatial location i, z_i*k*_ is the standardized value of ecological program *k*, and w*_k_* represents a predefined biological weight reflecting whether a given ecological program was considered to increase or decrease ecological opportunity. Programs associated with primitive leukemia, hypoxia, stromal support, extracellular matrix remodeling, immune suppression, and inflammatory remodeling were assigned positive weights (w*_k_* = +1), whereas programs associated with immune-active or potentially tumor-restrictive states were assigned negative weights (w*_k_* = −1). The resulting opportunity score is therefore an interpretable composite index summarizing the balance between ecological programs hypothesized to promote versus restrict leukemia persistence within the tissue microenvironment. Opportunity scores were subsequently projected onto tissue coordinates to generate spatial ecological opportunity landscapes.

### Spatial ecological context discovery

Spatial ecological contexts were identified jointly across the analyzed GSE279576 Visium sections. For each spatial capture location, 17 prespecified ecological program scores were assembled:

AML blast − like, primitive − like AML, committed − like AML, HSPC primitive, myeloid/ granulocytic, monocyte/macrophage, inflammatory, *CXCL*12 − *CXCR*4 axis, MSC/stromal, endothelial/vascular, osteolineage/endosteal, adipocytic, ECM/matrix, hypoxia/stress, T/NK, B − cell, and immune − suppression.

Only ecological-score columns available in the pooled dataset were retained. Nonfinite values were converted to missing values, and missing values were replaced with zero before standardization. The resulting ecological feature matrix was standardized across all pooled spatial spots using scikit-learn’s StandardScaler, which centers each feature at its pooled mean and scales it to unit variance:

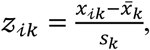

where *x*_i*k*_ is the ecological program score for spot i and feature *k*, and *x̅_k_* and *s_k_* are the corresponding mean and standard deviation calculated across the combined spatial dataset. Standardization was therefore performed jointly rather than independently within individual tissue sections.

Principal-component analysis was applied to the standardized ecological feature matrix using scikit-learn. The number of retained components was set to the smaller of eight and the number of available ecological features. The first five ecological principal components, EcoPC1–EcoPC5, were used as the input representation for clustering. The first two components were additionally used to visualize the resulting contexts in two-dimensional ecological space.

Unsupervised clustering was performed using the scikit-learn implementation of *k*-means with five clusters:

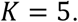

The algorithm minimized the within-cluster sum of squared Euclidean distances in the five-dimensional ecological principal-component space,

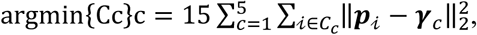

where ***p***_i_ denotes the EcoPC1–EcoPC5 coordinate vector for spot i, *C_c_* denotes cluster *c*, and ***γ****_c_* is the corresponding cluster centroid. Clustering used *k*-means++ centroid initialization, 50 independent initializations, and a fixed random seed of 0:

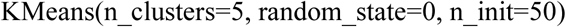

The solution with the lowest within-cluster sum of squares among the 50 initializations was retained. The five-context solution was specified a priori for the principal analysis; alternative values of *K* were not formally compared using silhouette scores, consensus clustering, or other cluster-number selection criteria. The resulting groups were therefore treated as candidate ecological contexts used to construct an interpretable intermediate representation rather than as uniquely determined biological states.

Because raw *k*-means cluster identifiers are arbitrary, clusters were relabeled S1–S5 using a predefined ordering score calculated from their mean ecological program values:

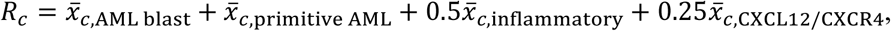

where *x̅_c_*_,*k*_ denotes the mean unstandardized ecological program score for cluster *c*. Raw clusters were ranked in descending order of *R_c_* and assigned sequentially as S1, S2, S3, S4, and S5. Thus, the S1–S5 numbering reflects an ordering based primarily on AML blast-like and primitive-like programs, with smaller contributions from inflammatory and *CXCL12*–*CXCR4*-axis programs, rather than the arbitrary numerical labels returned by *k*-means.

For biological interpretation, the mean value of each ecological feature was calculated within each context:

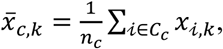

where *n_c_* is the number of spots assigned to context *c*. These context-level feature means were standardized across the five contexts for heatmap visualization. Candidate context annotations were assigned according to the ecological programs most strongly enriched within each group and their relative representation across bone marrow and extramedullary tissue sites. These annotations describe composite ecological-program patterns and were not interpreted as definitive cell-type identities.

Context composition within each sample was calculated as

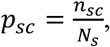

where *n_sc_* is the number of spots assigned to context *c* in sample *s*, and *N_s_* is the total number of analyzed spots in that sample. Site-level compositions were calculated analogously after grouping samples as bone marrow or extramedullary. Finally, context assignments were mapped back to the corresponding AnnData objects by spot barcode and projected onto the original Visium tissue coordinates for spatial visualization.

### Evolutionary opportunity field

To estimate context-level evolutionary opportunity, we summarized each ecological context using its standardized ecological feature profile. Contexts were embedded in ecological space and connected by pairwise similarity or neighborhood relationships. An opportunity score was assigned to each context based on malignant, primitive, hypoxic, stromal-supportive, immune-suppressed, and inflammatory features. Context-level opportunity scores were then projected onto spatial spots, generating tissue-level opportunity fields. The complete list of ecological features, assigned weights, signs, and component contributions is provided in Supplementary Data 6.

### OU-like ecological attractor summaries

OU-like summaries were used to characterize context-specific organization of spatial ecological states. These summaries were constructed from a one-dimensional latent ecological-state axis and a spatially ordered autocorrelation approximation rather than by fitting a continuous-time temporal Ornstein–Uhlenbeck process.

To construct the latent ecological-state variable, we selected eight ecological programs representing malignant, primitive hematopoietic, myeloid, monocyte–macrophage, inflammatory, and hypoxia-associated biology:

AML blast − like, primitive − like AML, committed − like AML, HSPC primitive, myeloid⁄granulocytic, monocyte⁄macrophage, inflammatory, hypoxia/stress.

For each spatial spot i, the selected program scores were standardized across the combined spatial dataset using feature-wise z-score transformation. Principal-component analysis was then applied to the standardized feature matrix, and the first principal component was retained as the latent ecological-state value *x*_i_. Because principal-component orientation is arbitrary, the sign of PC1 was oriented so that larger values corresponded to greater AML/primitive-like burden. Specifically, the mean of the AML blast-like, primitive-like AML, committed-like AML, and HSPC-primitive scores was used as an anchor. PC1 was multiplied by −1 when its correlation with this anchor was negative.

For each sample *s* and ecological context *c*, the context-specific attractor position was defined as the mean latent ecological state,

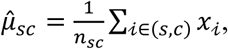

where *n_sc_* is the number of spots assigned to context *c* in sample *s*. The corresponding latent-state dispersion was summarized by

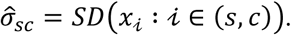

To obtain a spatial ordering within each sample–context combination, the two-dimensional tissue coordinates were centered and subjected to principal-component analysis. The first coordinate-derived principal component was used as a one-dimensional spatial pseudo-order. Spots were sorted according to this pseudo-order, producing an ordered latent-state sequence

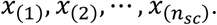

For groups containing at least six ordered spots, the lag-1 spatial correlation was calculated as

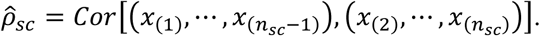

An AR(1)-like spatial decorrelation index was then defined as

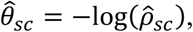

provided that

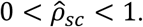

Values of 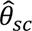 were left undefined when the group contained fewer than six spots, when either lagged sequence had zero variance, or when the estimated lag-1 correlation was nonpositive or greater than or equal to one. For visualization, positive spatial rate-index estimates were transformed as

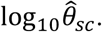

Under this parameterization, 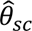 is inversely related to the persistence of neighboring latent-state values along the spatial pseudo-order. Values near zero arise when 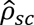 approaches one and indicate slowly changing or strongly persistent spatial organization. Larger 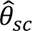 values arise from weaker positive lag-1 correlation and indicate more rapid spatial decorrelation. Accordingly, 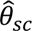 is best interpreted as an effective spatial decorrelation or restoring-rate index, rather than simply as a direct measure of retention strength.

Context-level parameter summaries were generated across samples for S1–S5. For spatial visualization, each spot was assigned the 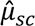, 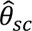, and log_10_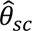 values corresponding to its sample and ecological context. Figure 4 used GSM8576301_BM1 as the representative spatial section.

Approximate attractor-basin curves were constructed for each ecological context using

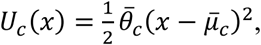

where 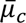 and 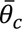 are the arithmetic means of the available sample-specific estimates for context *c*. Each curve was shifted vertically so that its minimum was zero. These curves provide qualitative visualizations of context-specific latent-state centers and curvature and were not obtained from likelihood-based fitting of a temporal stochastic differential equation.

Because the datasets consist of cross-sectional tissue sections, the resulting quantities do not represent longitudinal OU equilibrium positions or temporal mean-reversion rates. They are operational spatial summaries derived from latent ecological-state distributions and autocorrelation along a tissue-coordinate-based pseudo-order.

### Lévy-like ecological escape analysis

Lévy-like ecological escape was operationally defined as cross-context spatial adjacency combined with ecological separation between the corresponding context profiles. The analysis was performed separately within each GSE279576 tissue section so that spatial neighbor relationships were never constructed across samples.

For each sample, spots with available tissue coordinates and ecological-context assignments were represented by their two-dimensional Visium coordinates,

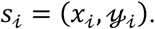

A spatial *k*-nearest-neighbor graph was constructed using Euclidean distance in tissue-coordinate space with

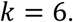

The scikit-learn NearestNeighbors implementation was fitted separately to each sample using n_neighbors=7, including the spot itself, and the six nonself nearest neighbors were retained. Because nearest-neighbor relationships can be returned in both directions, each unordered spot pair was included only once. Thus, the resulting graph comprised unique undirected spatial-neighbor edges within each tissue section.

Each neighbor edge ℯ = (i, j) was classified as either a same-context edge or a cross-context edge according to the ecological-context assignments *c*_i_ and *c*_j_:

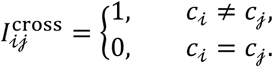

Ecological separation among contexts was calculated from the standardized context-level ecological feature profiles. Let

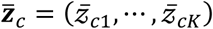

denote the standardized ecological feature vector for context *c*. The ecological jump distance between two contexts *a* and *b* was defined as their Euclidean distance:

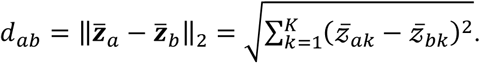

Every spatial-neighbor edge connecting contexts *a* and *b* was assigned the corresponding context-level ecological distance *d_ab_*. Therefore, all edges connecting the same pair of ecological contexts had the same ecological jump distance, although their physical locations and spatial distances differed.

For descriptive purposes, the physical Euclidean distance between neighboring spots was also retained. When the latent ecological-state value ℓ_i_ was available, the absolute latent-state displacement was calculated as

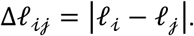

This latent-state displacement was stored in the edge table but was not used to define the primary Lévy-like escape score.

Rare ecological jumps were identified from the empirical distribution of ecological distances among all cross-context neighbor edges. The rare-jump threshold was defined as the 90th percentile:

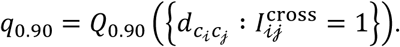

A spatial-neighbor edge was classified as a rare jump when it connected two different contexts, and its ecological distance was greater than or equal to this threshold:

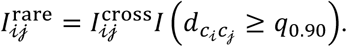

The threshold was estimated from the pooled cross-context edge distribution across the analyzed tissue sections. Rare-jump edges were subsequently projected onto the original Visium coordinates to identify their spatial locations within representative sections.

For context-level scoring, each undirected neighbor edge was expanded into two directed records so that both endpoint contexts could be evaluated as source contexts. For ecological context *c*, let ℰ*_c_* denote all directed spatial-neighbor edges originating from that context. The cross-context transition frequency was calculated as

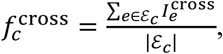

and the rare-jump frequency was calculated as

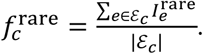

The mean ecological distance among outgoing cross-context edges was

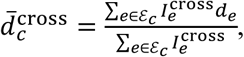

with a value of zero assigned when no outgoing cross-context edges were present. Similarly, the mean distance among rare outgoing edges was

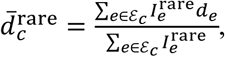

with zero assigned when no rare edges were present.

The primary context-level Lévy-like escape score was defined as

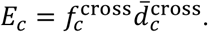

Thus, a context received a high escape score when a large proportion of its local spatial-neighbor relationships crossed into other ecological contexts and those transitions connected ecologically distant context profiles. A complementary rare-escape score was calculated as

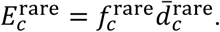

The primary score therefore summarizes the frequency-weighted ecological magnitude of all cross-context transitions, whereas the rare-escape score isolates transitions in the upper 10% of the cross-context ecological-distance distribution.

These quantities were termed Lévy-like because they distinguish frequent local retention or short ecological transitions from comparatively uncommon, large cross-context displacements. They do not establish that the data were generated by a formal Lévy stochastic process. Demonstrating a generative Lévy process would require explicit distributional fitting, comparison with alternative jump models, and preferably longitudinal spatial observations.

### Branching-like amplification analysis

Branching-like amplification was defined as an interpretable composite measure that integrates ecological abundance, latent-state displacement, ecological opportunity, Lévy-like escape, and the proportion of escaped states. For each ecological state i, we calculated a branching amplification score,

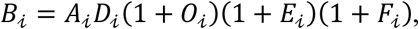

where

- *B*_i_ denotes the branching-like amplification score;
- *A*_i_ is the relative abundance of ecological state i;
- *D*_i_ is the latent-state displacement from the corresponding resident ecological state, representing the magnitude of ecological divergence;
- *O*_i_ is the evolutionary opportunity score;
- *E*_i_ is the Lévy-like escape score summarizing the frequency-weighted ecological magnitude of cross-context neighbor transitions; and
- *F*_i_ is the escaped-state fraction, defined as the proportion of observations assigned to escaped ecological states.

The multiplicative formulation was chosen to represent the hypothesis that ecological amplification emerges from the combined influence of population abundance, ecological divergence, favorable opportunity, rare ecological escape, and the accumulation of escaped states. The additive offsets (1 + *O*_i_, 1 + *E*_i_, and 1 + *F*_i_) preserve positive scaling while allowing each component to contribute proportionally to the composite score. For the treatment-associated branching analysis, opportunity was represented by the available state-level opportunity variable. When no independently estimated opportunity value was available in the GSE235923-derived validation dataset, *O*_i_ was fixed at 1.0 for all states. Consequently, the factor (1 + *O*_i_) = 2 was constant and did not contribute to differences in branching-amplification scores within this analysis.

Ecological states with branching amplification scores in the upper quartile of the empirical distribution were classified as amplified states. Resident, escaped, and amplified states were visualized using a low-dimensional embedding of the latent ecological representation. A state-amplification network was then constructed by connecting resident or low-escape states to high-scoring escaped or amplified states, with node size proportional to relative abundance (*A*_i_) and edge width proportional to the corresponding branching amplification score (*B*_i_).

Because no lineage tracing or phylogenetic information is available in the analyzed datasets, the branching amplification score should be interpreted as an operational summary of ecological expansion potential rather than a direct estimate of cell proliferation, clonal fitness, or a branching-process birth rate. Instead, it provides an interpretable index that integrates multiple ecological properties associated with the spatial prominence and potential expansion of candidate tissue states.

### Therapy-response validation

Therapy-response validation was performed using diagnosis-sample ecological composition summaries derived from the GSE235923 secondary cohort. For each sample *s*, a primitive-state retention proxy was defined as the combined relative abundance of HSC-like and progenitor-like states:

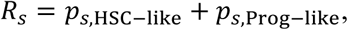

where *p_s_*_,HSC−like_ and *p_s_*_,Prog−like_ denote the fractions of cells assigned to the HSC-like and progenitor-like ecological states, respectively. Because the four principal state fractions were normalized within each sample, *R_s_* represents the proportion of the principal ecological composition occupying primitive or progenitor-associated states.

This sample-level compositional proxy is distinct from the spatial OU-like quantities estimated from GSE279576 in Figure 4. Specifically, it was not calculated from the lag-1 spatial autocorrelation *ρ*, the spatial rate index *θ* = −log(*ρ*), or any transformation of those parameters. The term retention is used here operationally to denote preservation of primitive HSC/progenitor-associated composition in the treatment-associated validation cohort.

For each sample, the primitive-state retention proxy was analyzed together with the Lévy-like escape score and branching-like amplification score. Each component was standardized across samples using a z-score, and the three standardized values were summed:

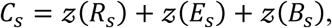

where *E_s_* is the sample-level Lévy-like escape score and *B_s_* is the branching-like amplification score. The resulting quantity was interpreted as an integrated ecological composite score rather than a validated clinical-risk score.

Component scores were compared between relapse-associated and other/unknown samples using two-sided Mann–Whitney U tests. Tests were exploratory because the groups contained *n* = 2 relapse-associated and *n* = 17 other/unknown samples. Exact unadjusted *P* values were reported in Figure 6E, and no multiple-testing adjustment was applied.

### Software availability and reproducibility

Figures were generated using Python and refined in Adobe Illustrator for publication-quality layout. Python analyses used pandas, numpy, scanpy, scikit-learn, scipy, matplotlib, seaborn, and networkx. Final outputs included figure panels, summary tables, context annotations, opportunity scores, OU-like parameter summaries, Lévy-like escape scores, branching amplification summaries, therapy-response summaries, and network edge tables. All executable scripts, processed intermediate outputs, environment specifications, and figure-generation workflows are publicly available at the project GitHub repository and are archived as a versioned release through Zenodo at doi:10.5281/zenodo.21633622.

## DATA ACCESS

Public datasets used in this study were obtained from the Gene Expression Omnibus (GEO) under accession numbers GSE253355, GSE279576, and GSE235923.

The complete computational workflow, including data preprocessing, ecological program scoring, spatial ecological context discovery, evolutionary opportunity scoring, OU-like ecological retention analysis, Lévy-like ecological escape analysis, branching-like ecological amplification, therapy-response validation, and figure generation, is publicly available through the project GitHub repository: https://github.com/shkim9391/Spatial_Therapy_OU_Levy_Branching

The repository includes executable analysis scripts, software environment specifications, processed intermediate and summary tables, figure-generation workflows, and documentation linking each computational step to the corresponding manuscript figure. An archived, versioned snapshot of the complete workflow and processed outputs is permanently available through Zenodo: https://doi.org/10.5281/zenodo.21633622

No new sequencing data were generated for this study.

## ACKNOWLEDGMENTS

This study was conducted using publicly available genomic and spatial transcriptomic datasets generated by the original investigators, whose contributions are gratefully acknowledged. The author thanks the research communities responsible for producing and maintaining these valuable public resources, which made this work possible.

The author also acknowledges the support of the Department of Pediatric Oncology at Dana-Farber Cancer Institute for providing a stimulating scientific environment during the development of this work. Computational analyses were performed using open-source software developed by the scientific Python community, including Scanpy, NumPy, pandas, SciPy, scikit-learn, and Matplotlib.

The conclusions presented in this study are solely those of the author and do not necessarily reflect the views of the affiliated institutions.

## Author Contributions

S.-H.K. conceived the study, designed the computational framework, performed data preprocessing and statistical analyses, developed the ecological modeling methodology, generated all figures, interpreted the results, wrote the manuscript, and approved the final version for publication.

## Supplemental Material

Supplemental Data 1–10 provide the dataset inventory, ecological program definitions, context profiles and annotations, opportunity-score components, ecological transition summaries, OU-like parameter estimates, Lévy-like escape summaries, and branching and therapy-validation outputs.

